# Translational profiling of macrophages infected with *Leishmania donovani* identifies mTOR- and eIF4A-sensitive immune-related transcripts

**DOI:** 10.1101/2019.12.20.884338

**Authors:** Visnu Chaparro, Louis-Philippe Leroux, Laia Masvidal, Julie Lorent, Tyson E. Graber, Aude Zimmermann, Guillermo Arango Duque, Albert Descoteaux, Tommy Alain, Ola Larsson, Maritza Jaramillo

## Abstract

The protozoan parasite *Leishmania donovani* (*L. donovani*) causes visceral leishmaniasis, a chronic infection which is fatal when untreated. While previous studies showed that *L. donovani* reprograms transcription to subvert host cell functions, it remains unclear whether the parasite also alters host mRNA translation to establish a successful infection. To assess this, we compared transcriptome-wide translation in primary mouse macrophages infected with *L. donovani* promastigotes or amastigotes using polysome-profiling. This identified ample selective changes in translation (3,127 transcripts) which were predicted to target central cellular functions by inducing synthesis of proteins related to chromatin remodeling and RNA metabolism while inhibiting those related to intracellular trafficking and antigen presentation. Parallel quantification of protein and mRNA levels for a set of transcripts whose translation was activated upon *L. donovani* infection (*Papbpc1, Eif2ak2,* and *Tgfb*) confirmed, as indicated by polysome-profiling, increased protein levels despite largely unaltered mRNA levels. Mechanistic *in silico* analyses suggested activated translation depending on the kinase mTOR (e.g. *Pabpc1*) and the RNA helicase eIF4A (e.g. *Tgfb*) during infection. Accordingly, treatment with mTOR inhibitors torin-1 or rapamycin reversed *L. donovani*-induced PABPC1 without affecting corresponding transcript levels. Similarly, the production of TGF-β decreased in presence of the eIF4A inhibitor silvestrol despite unaltered *Tgfb* mRNA levels. Consistent with parasite modulation of host eIF4A-sensitive translation to promote infection, silvestrol suppressed *L. donovani* replication within macrophages. In contrast, parasite survival was favored under mTOR inhibition. In summary, infection-associated changes in translation of mTOR- and eIF4A-sensitive mRNAs contribute to modulate mRNA metabolism and immune responses in *L. donovani*-infected macrophages. Although the net outcome of such translation programs favours parasite propagation, individual translation programs appear to have opposing roles during *L. donovani* infection, thereby suggesting their selective targeting as key for therapeutic effects.

**Author Summary:** Fine-tuning the efficiency of mRNA translation into proteins allows cells to tailor their responses to stress without the need for synthesizing new mRNA molecules. It is well established that the protozoan parasite *Leishmania donovani* alters transcription of specific genes to subvert host cell functions. However, discrepancies between transcriptomic and proteomic data suggest that post-transcriptional regulatory mechanisms also contribute to modulate host gene expression programs during *L. donovani* infection. Herein, we report that one third of protein-coding mRNAs expressed in macrophages are differentially translated upon infection with *L. donovani*. Our computational analyses reveal that subsets of mRNAs encoding functionally related proteins share the same directionality of translational regulation, which is likely to impact metabolic and microbicidal activity of infected cells. We also show that upregulated translation of transcripts that encode central regulators of mRNA metabolism and inflammation is sensitive to the activation of mTOR or eIF4A during infection. Finally, we observe that inhibition of eIF4A activity reduces parasite survival within macrophages while selective blockade of mTOR has the opposite effect. Thus, our study points to a dual role for translational control of host gene expression during *L. donovani* infection and suggests that novel regulatory nodes could be targeted for therapeutic intervention.

## Introduction

Visceral leishmaniasis (VL) is a vector-borne infection caused by protozoan parasites of the *Leishmania donovani* (*L. donovani*) complex. VL is endemic in more than 60 countries and is frequently lethal if untreated (1). The lack of efficient vaccines and the failure to control emerging parasite resistance reflect the urgent need to design safe and efficient therapeutics targeting host-encoded factors (2). In mammalian hosts, *Leishmania* promastigotes preferentially colonize macrophages, where they transform into replicative amastigotes that proliferate within modified phagolysosomes (1). To establish a successful infection, the parasite dampens antimicrobial responses, alters vesicle trafficking, and subverts immunomodulatory functions and metabolic processes of the host cell (3).

At the molecular level, *L. donovani* modulates the activity of multiple host cell signaling pathways and transcription factors (3). Consistently, profiling of mRNA levels in *L. donovani*-infected macrophages revealed vast perturbation in host gene expression programs associated with parasite persistence (4–8). The pioneering data supporting widespread changes in host cell mRNA levels following *L. donovani* infection were obtained in bone marrow-derived macrophages (BMDMs) using cDNA-microarrays (4). This study showed that *L. donovani* axenic amastigotes downregulate expression of genes involved in apoptosis and NF-κB signaling while stimulating those encoding monocyte chemoattractants. Subsequently, DNA-microarray based studies of human and mouse monocyte-derived macrophages infected with *L. donovani* promastigotes identified increased levels of transcripts related to cell migration and repression of genes encoding MHC class II molecules (5, 6). More recently, RNA sequencing (RNAseq) of mouse peritoneal macrophages infected with *L. donovani* showed a strong suppression of genes related to immune activation, signal transduction, phagosome, and endocytosis (8). Remarkably, combined analysis of RNAseq data from cells and tissues of infected hamsters provided evidence that despite a strong pro-inflammatory signature in the spleen, *L. donovani* induced a complex gene expression pattern in splenic macrophages characterized by M1- and M2-associated transcripts that skews their responses to IFN-γ, thereby rendering them more susceptible to the infection (7). Altogether, these studies support an important role of parasite-directed reprogramming of the host transcriptome in the immunopathogenesis of VL. However, discrepancies between transcriptomics and proteomics data of *L. donovani*-infected macrophages (9) suggest that post-transcriptional and post-translational mechanisms may also modulate the host cell proteome during VL infection.

Among post-transcriptional mechanisms, transcript-selective changes in translational efficiencies enable cells to swiftly remodel their proteomes in response to environmental cues without requiring *de novo* mRNA synthesis (10–13). In eukaryotes, translational efficiency is mainly regulated at the initiation step when ribosomes are recruited to the mRNA (14). This process is facilitated by the eukaryotic translation initiation factor 4F (eIF4F) complex, consisting of eIF4E, the mRNA 5’-m7G-cap-binding subunit; eIF4G, a scaffolding protein; and eIF4A, an RNA helicase (14). Activation of the mechanistic target of rapamycin (mTOR) complex 1 stimulates formation of the eIF4F complex which promotes translation of mRNAs that are particularly sensitive to changes in eIF4E levels and/or availability. These include those containing a 5’ terminal oligopyrimidine (5’ TOP) motif (15, 16), those with highly structured 5’ UTR sequences, which are largely dependent on the RNA helicase activity of eIF4A for their translation (17), and those with very short 5’UTRs (16, 18). Many mTOR- and eIF4A-sensitive mRNAs encode proteins related to translation, cell survival, metabolism, proliferation, and growth (16, 18, 19). Interestingly, a number of innate immune regulators are also under translational control via mTOR- or eIF4A-dependent mechanisms (20, 21), including several pro- and anti-inflammatory mediators in macrophages (11–13, 22). Thus, key immune cell functions may be hijacked by intracellular pathogens via modulation of mRNA translation. Here we show that infection with promastigotes or amastigotes of *L. donovani* leads to an early translational reprogramming in macrophages partially depending on mTOR and eIF4A activity which appears to contribute to both parasite persistence and host cell defense.

## Results

### *L. donovani* selectively modulates the macrophage translatome

To investigate the impact of *L. donovani* infection on the host cell translatome (i.e. the pool of efficiently translated mRNAs) at a transcriptome-wide level, BMDMs were incubated with *L. donovani* promastigotes or amastigotes for 6 h and compared to uninfected cells using polysome-profiling quantified by RNAseq (**Fig 1A**). Polysome-profiling determines levels of both efficiently translated mRNA and cytoplasmic mRNA. Such data enables identification of bona fide changes in translation efficiencies (i.e. changes in levels of polysome-associated mRNA which are not paralleled by corresponding changes in cytoplasmic mRNA levels) using the anota2seq algorithm (23). At a false discovery rate (FDR) ≤0.15, anota2seq identified widespread mRNA-selective alterations in translational efficiencies upon infection with either parasite life stage (**Fig 1B**). From a total of 9,442 host protein-encoding mRNAs detected, 27% showed altered translational efficiency following infection with *L. donovani* amastigotes (13% increased and 14% reduced) (**Fig 1C**, left panel; **Fig 1D** top panels, and **S1 Table**). Similarly, the translational efficiency of 18% of the host cell transcripts was altered in response to *L. donovani* promastigotes (9% increased and 9% decreased) (**Fig 1C**, right panel; **Fig 1D**, bottom panels, and **S1 Table**). Consistent with changes in translational efficiencies largely independent of parasite stage, only 1.5% of all mRNAs differed between host cells infected with *L. donovani* promastigotes as compared to amastigotes (**S1 Fig** and **S1 Table**). In addition to detecting differences in translational efficiency affecting protein levels, anota2seq allows for identification of transcripts whose changes in mRNA levels are buffered at the level of translation such that their polysome-association remains largely unaltered (23). This is a mode for regulation of gene expression which offsets the relationship between mRNA levels and protein levels to suppress changes in protein levels imposed by altered transcription or mRNA stability (24). Interestingly, a large subset of transcripts whose abundance changed upon infection with *L. donovani* amastigotes or promastigotes was buffered at the level of translation (21% out of 1,051 and 29% out of 1,604 mRNAs, respectively) (**Fig 1D** and **S1 Table**). In contrast, only a small number of transcripts (71 mRNAs) whose levels differed between *L. donovani* promastigote- and amastigote-infected BMDMs (**S1B-C Figs**) were translationally buffered. These data indicate that both life stages of *L. donovani* induce abundant and largely similar selective changes in translational efficiency of host cell mRNAs that modulate or maintain protein levels.

**Figure 1.**
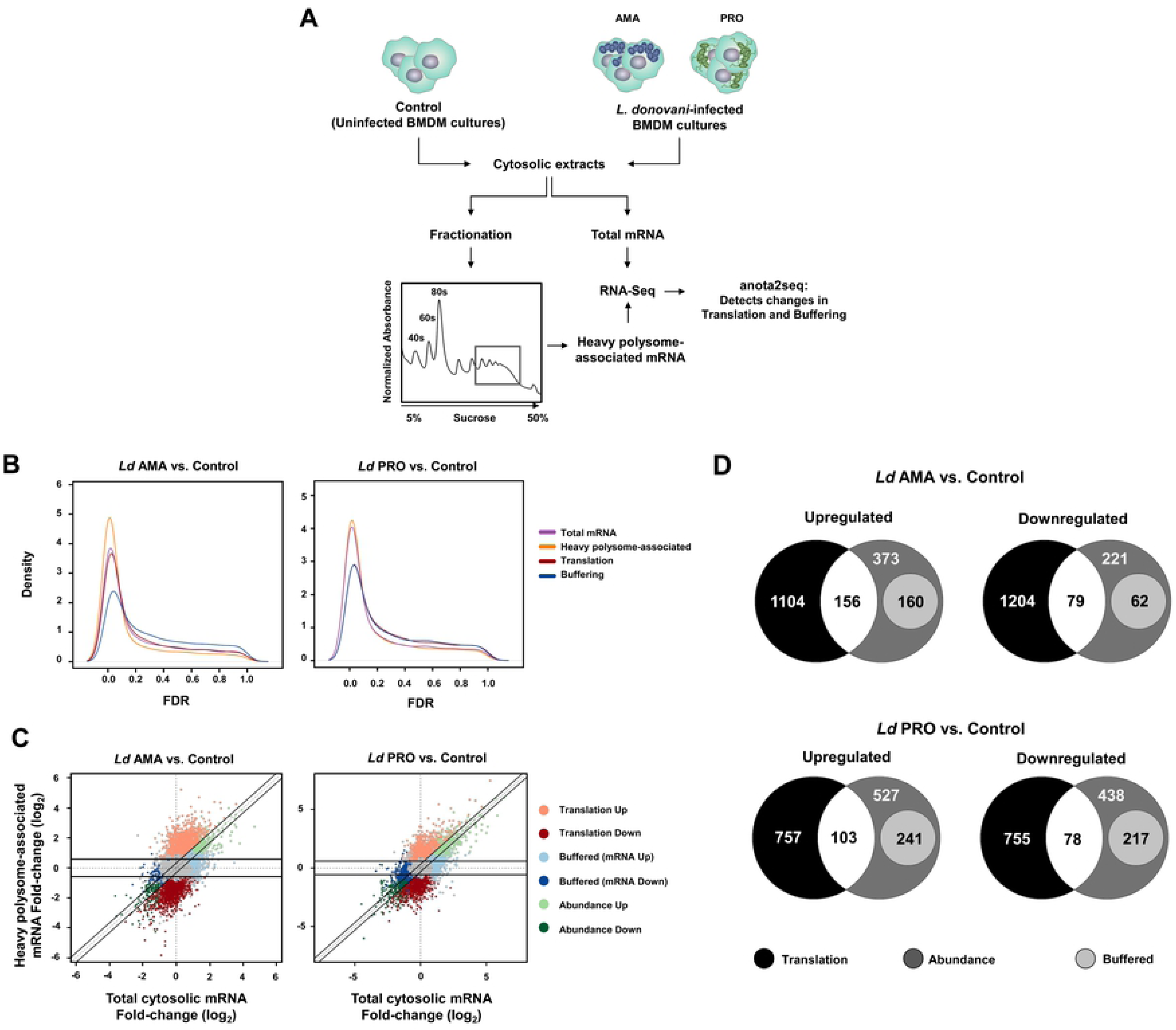
*L. donovani* infection causes widespread changes in mRNA translation in macrophages. (**A**) Strategy to characterize the translatome of *L. donovani*-infected BMDMs. (**B**) Kernel densities of adjusted *p* values (FDRs) following anota2seq analysis of changes in total mRNA, heavy polysome-associated mRNA, translation (i.e. changes in polysome-associated mRNA not paralleled by changes in total mRNA) and translational buffering (i.e. changes in total mRNA buffered at the level of translation) for BMDMs infected with *L. donovani* amastigotes (*Ld* AMA) or promastigotes (*Ld* PRO) compared to uninfected cells (Control). (**C**) Scatter plots of log_2_ fold changes (for the same comparisons as in panel B) for heavy polysome-associated mRNA and total cytosolic mRNA. Differentially regulated transcripts through translation, abundance or buffering are indicated. Unchanged mRNAs are shown in grey. (**D**) Venn diagrams showing the number of mRNAs up- or down-regulated at the level of translation, abundance, and buffering for BMDMs infected with *Ld* AMA or *Ld* PRO compared to Control. (**B-D**) Data analyses were performed on samples generated from at least three biological replicates.

### Transcript-selective changes in translation upon *L. donovani* infection target a variety of macrophage functions

To assess how the host cell phenotype is potentially affected by mRNA-selective perturbation of translation during *L. donovani* infection, we searched for enrichment of cellular functions defined by Gene Ontology (GO) classifications (25) among proteins encoded by transcripts showing altered translational efficiencies (**Fig 2A** and **S2 Table**). Enriched categories, among proteins encoded by translationally activated transcripts in BMDMs infected by either parasite life stage, included chromatin remodeling, regulation of mRNA metabolism (i.e. splicing, export from the nucleus, stability and translation), regulation of type I IFN production and protein deubiquitination (**Fig 2A**, left panel; FDR ≤0.05). Accordingly, mRNAs encoding histone modifying enzymes (e.g. *Ash1l*, *Ep300*, *Kmt2a*, *Kmt4c*), transcription factors (e.g. *Cebpb*, *Foxo4*, *Ets2*, *Elk4*), translation initiation factors (e.g. *Eif3a/b/c*, *Eif4g3*), ribosomal proteins (e.g. *Rpl13a*, *Rpl38*, *Rps12*), RNA-binding proteins (RBPs) (e.g. *Pabpc1*, *Eif2ak2*, *Larp1*, *Pum1*), and ubiquitin hydrolases (e.g. *Usp25*, *Fam63b, Usp36*) showed increased translational efficiency in *L. donovani*-infected macrophages compared to uninfected controls (log_2_ fold-change ˃1.0, FDR ≤0.15) (**Fig 2B**, top panels). In contrast, proteins encoded by mRNAs whose translational efficiency was suppressed upon infection by either life stage of *L. donovani* were enriched for categories related to protein trafficking (i.e. Rab protein signal transduction, vesicle organization, and post-Golgi vesicle-mediated transport), cell metabolism (i.e. mitochondrial membrane organization, mitochondrial respiratory chain complex assembly, fatty acid beta-oxidation, and peroxisomal membrane transport), protein ubiquitination, and tRNA metabolism (**Fig 2A**, right panel; FDR ≤0.05). Specifically, mRNAs translationally suppressed by promastigotes or amastigotes of *L. donovani* encode proteins involved in antigen presentation (e.g. *Cd74*, *H2-Q1*), intracellular transport (e.g. *Rab18*, *Sec22a*, *Vamp3*, *Vps37c*), organization of lysosome (e.g. *Bloc1s2*, *Laptm4b*), mitochondria (e.g. *Cox18*, *Ndufa8*, *Timm21*, *Tomm22*), peroxisome (e.g. *Paox*, *Pex2*, *Pex7*) and Golgi apparatus (e.g. *Golga7*, *Tango2*), lipid metabolism (e.g. *Nr1h2*, *Pla2g6*, *Scp2*), protein ubiquitination (e.g. *Trim68*, *Ube2g1*, *Ube2w*), and tRNA modifications (e.g. *Trmo*, *Tsen15, Trmt12*) (log_2_ fold-change < −1.0, FDR ≤0.15) (**Fig 2B**, bottom panels). Interestingly, several mRNAs encoding innate immune sensors showed activated translation (e.g. *Dhx9*, *Tlr9*, and *Zbtb20*) while others were translationally suppressed (e.g. Clec7a, *Mavs*, *Tlr2*), possibly suggesting a dual role for translation-dependent alterations in macrophage immune functions during *L. donovani* infection.

**Figure 2.**
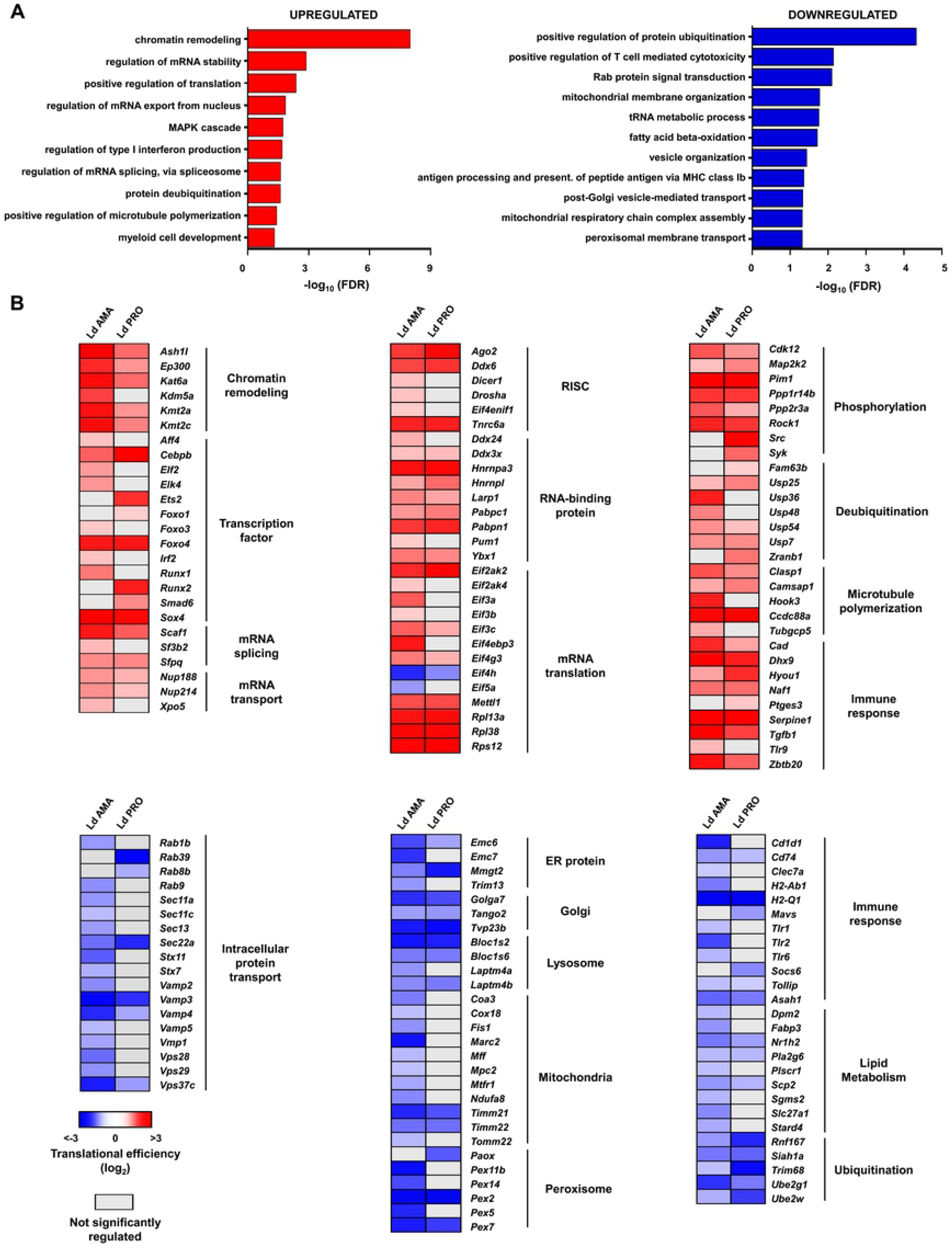
*L. donovani* infection-dependent translation targets core and immune cell functions. (**A**) FDR values (-log_10_) for selected GO term enriched categories for translationally up- and down-regulated host mRNAs upon *L. donovani* infection. (**B**) Heatmaps showing changes in translational efficiencies for selected genes in enriched GO terms. Analyses were carried out on data generated from at least three biological replicates.

### *L. donovani* infection enhances mTOR-sensitive mRNA translation in macrophages

We next sought to identify upstream mechanisms underlying observed changes in selective mRNA translation. Activation of Akt and ribosomal protein kinase 1 (S6K1) has been reported in macrophages as early as 30 min post-infection with *L. donovani* promastigotes (26). As altered activity of the PI3K/AKT/mTOR pathway modulates mRNA translation in a selective fashion (16, 18, 27), we characterized the kinetics of mTOR activity upon *L. donovani* infection. To this end, we monitored the phosphorylation status of its downstream targets S6K1 and eIF4E-binding protein 1 (4E-BP1) in BMDMs. Transient phosphorylation of S6K1 at T389 was observed in BMDMs infected by either *L. donovani* promastigotes or amastigotes (**Fig 3A**). Accordingly, phosphorylation of S6K1 substrate ribosomal protein S6 (RPS6) at S235, S236, S240, and S244 was augmented during infection (**Fig 3A**). In addition, early phosphorylation of 4E-BP1 at T37/46 was induced with similar kinetics by both parasite life stages (**Fig 3A**). To assess whether mTOR-sensitive translation was regulated in *L. donovani*-infected BMDMs, we focused on 5’ TOP-containing mRNAs, whose translation is highly dependent on mTOR activity and encode for ribosomal proteins and translation initiation and elongation factors (16, 18, 27). Indeed, translation of previously described TOP-mRNAs (27) was selectively activated during infection independent of parasite stage (*p* < 0.001 for both stages) (**Figs 3B-C**, and **S3A Table**). Thus, PI3K/AKT/mTOR signalling and downstream mTOR-sensitive translation are activated in BMDMs early during infection by *L. donovani* amastigotes or promastigotes.

**Figure 3.**
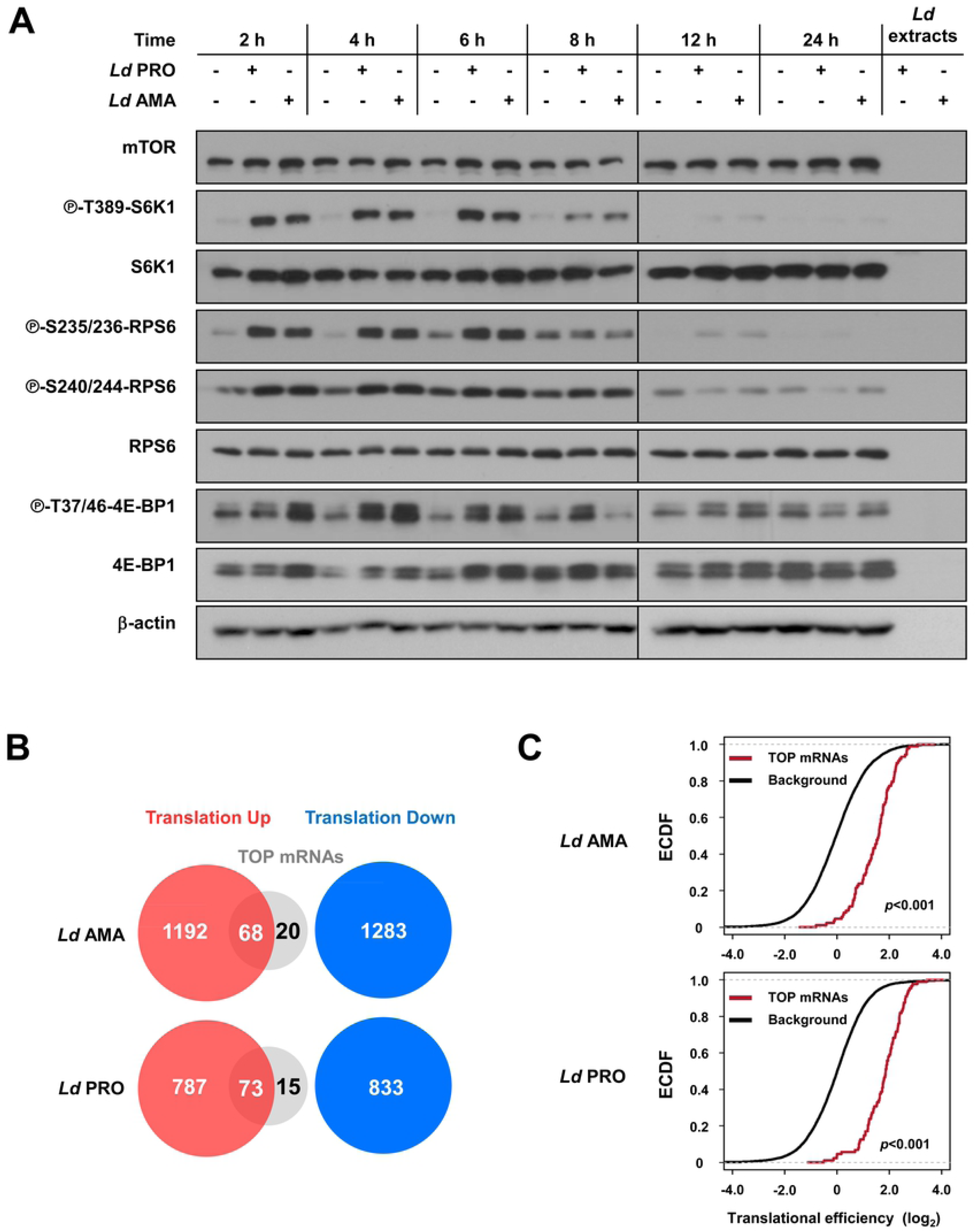
mTOR-sensitive host mRNA translation is activated during *L. donovani* infection. (A) BMDM cultures were inoculated with either *Ld* AMA or *Ld* PRO or left uninfected for the indicated times. Phosphorylation and expression levels of indicated proteins were monitored by Western blotting. Total amounts of β-actin were used as a loading control. Total protein extracts from *Ld* cultures were used to control for any cross-reactivity of the antibodies against parasite antigens. (**B**) Venn diagrams indicating the numbers of transcripts harboring a TOP-motif among translationally activated vs. suppressed transcripts following *Ld* AMA or *Ld* PRO infection as compared to control BMDMs. (**C**) Empirical cumulative distribution function (ECDF) of translational efficiencies for TOP mRNAs as compared to those of all detected transcripts (background). Differences in translational efficiencies between transcripts with TOP-motifs vs control transcripts were assessed per parasite stage. (**A**) Results are representative of at least three independent experiments. (**B**-**C**) Data analyses were performed on samples generated from at least three biological replicates.

### Translation of mRNAs encoding RNA-binding proteins PABPC1 and EIF2AK2 is activated during *L. donovani* infection in an mTOR-dependent fashion

A number of studies point to a central role for RBPs in coordinating macrophage inflammatory responses and anti-microbial activity (28), including poly(A)-binding protein cytoplasmic 1 (PABPC1) (29) and protein kinase dsRNA-activated (PKR, also known as EIF2AK2) (30, 31). Remarkably, anota2seq detected increased translational efficiency of *Pabpc1* and *Eif2ak2* mRNAs in BMDMs infected with *L. donovani* promastigotes or amastigotes (**Fig 2B**, top-middle panel and **S1 Table**). Accordingly, expression of both proteins increased upon infection (**Fig 4A**) without significant changes in mRNA abundance (**Fig 4B**). Moreover, phosphorylation of EIF2AK2 at T451 was augmented in response to infection with either parasite stage (**S2 Fig**). Surprisingly, phosphorylation of eIF2α at S51, a main downstream target of EIF2AK2 (32), decreased in *L. donovani*-infected BMDMs (**S2 Fig**). The *Pabpc1* mRNA was previously shown to contain a TOP-motif (15) and, consistently, BMDM treatment with mTOR inhibitors rapamycin or torin-1 suppressed PABPC1 protein expression during *L. donovani* infection (**Fig 4C**) independently of mRNA levels (**Fig 4D**, top panel). Of note, neither host cell (11) nor extracellular parasite viability (**S3 Fig**) were affected by mTOR inhibition. EIF2AK2 is induced by TLR stimuli such as LPS and poly (I:C) which also activate mTOR signaling (33, 34). Moreover, *L. donovani* activates TLR signaling (35). Interestingly, *L. donovani*-induced EIF2AK2 protein expression in BMDMs was reduced in presence of rapamycin or torin-1 (**Fig 4C**) independently of *Eif2ak2* mRNA levels (**Fig 4D**, bottom panel). In sum, this set of experiments indicates that *L. donovani* infection activates translation of *Pabpc1* and *Eif2ak2* mRNAs to increase PABPC1 and EIF2AK2 protein levels in BMDM by stimulating mTOR activity.

**Figure 4.**
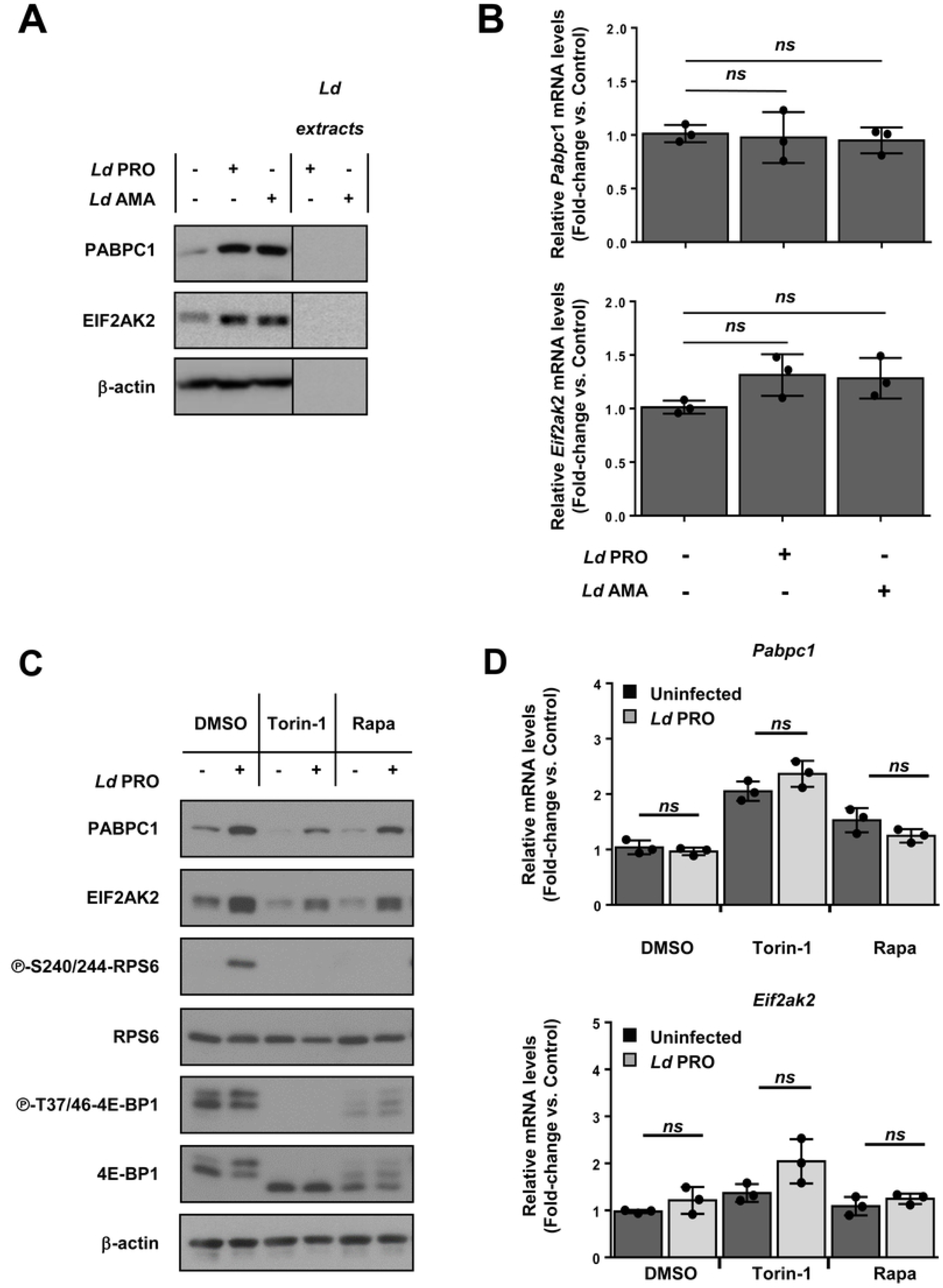
Upregulation of PABPC1 and EIF2AK2 in macrophages infected with *L. donovani* is mTOR-dependent. (**A**-**B**) BMDM cultures were inoculated with *Ld* AMA, *Ld* PRO or left uninfected for 6 h. (**C**-**D**) Cells were pre-treated with 200 nM of torin-1, 20 nM rapamycin (Rapa) or DMSO for 2 hours before infection. (**A**, **C**) Total levels of PABPC1 and EIF2AK2 were monitored by Western blotting. (**C**) Phosphorylation status of mTOR downstream targets RPS6 and 4E-BP1. (**B**, **D**) Relative mRNA amounts of *Pabpc1*and *Eif2ak2* (normalized to *Actb*) were measured by RT-qPCR. (**A**, **C**) Results are representative of three independent experiments. (**B**, **D**) Data are presented as mean ± SD (biological replicates, n=3). \*\**p* <0.01, \*\*\**p* <0.001 (for the indicated comparisons), *ns* = non-significant.

### Translation of eIF4A-sensitive mRNAs is activated upon *L. donovani* infection

As mentioned above, the RNA helicase eIF4A facilitates translation of transcripts harboring long and highly structured 5’ UTR sequences. Some such encoded proteins are involved in tumor immune evasion (36) and progression of viral (37) and protozoan parasitic infections (12), suggesting that eIF4A-dependent translation may contribute to herein observed changes in translational efficiencies (**Fig 1**). In addition to eIF4F-complex formation, the unwinding activity of eIF4A is enhanced by the translation initiation factor eIF4B (14). Consistent with eIF4B-dependent modulation of eIF4A activity, levels of phosphorylated and total eIF4B protein were increased in BMDMs infected with *L. donovani* amastigotes or promastigotes (**Fig 5A**). To test whether this may contribute to selective regulation of mRNA translation following parasite infection, we assessed translational efficiencies of a compilation of previously described eIF4A-sensitive transcripts (36, 38–40). Indeed, following infection independently of parasite stage, the translational efficiencies of such mRNAs were elevated as compared to background transcripts (*p* <0.001) (**Fig 5B**). From a total of 1198 previously described eIF4A-sensitive mRNAs, 149 were translationally activated whereas 80 were translationally suppressed upon infection with promastigotes or amastigotes of *L. donovani* (**S3B Table**). The presence of a 5’ UTR G-quadruplex-forming guanine quartet (CGG)_4_ motif is an indirect approach to assess whether transcripts are expected to be more dependent of eIF4A for their translation (41). Indeed, analysis of Motif Enrichment (AME) revealed a significant enrichment of the (CGG)_4_ motif in 5’ UTRs of transcripts with highly activated translation (≥4-fold increase in translational efficiency upon infection) as compared to 5’ UTRs from transcripts with unaltered translational efficiency (*p* = 0.0036) (**Fig 5C**). TGF-β is a key cytokine implicated in the distinctive immune suppression that follows *L. donovani* infection *in vivo* (42, 43). Upon infection with *L. donovani* amastigotes or promastigotes, anota2seq analysis identified augmented translational efficiency of the tumor growth factor-β 1 *(Tgfb1)* mRNA (**Fig. 2B**, top-right panel and **S1 Table**), which is highly dependent of eIF4A for its translation (39). Accordingly, production of TGF-β increased in BMDMs upon infection (**Fig 5D**) without changes in *Tgfb1* mRNA abundance (**Fig 5E**). Remarkably, inhibition of eIF4A activity using silvestrol abrogated TGF-β induction in *L. donovani*-infected BMDMs (**Fig 5D**) without affecting *Tgfb1* mRNA levels (**Fig 5E**). Of note, no acute toxicity was detected in BMDMs and extracellular parasites exposed to silvestrol (**S4 Fig**). Thus, these data indicate that, in addition to mTOR-dependent translation, *L. donovani* infection also bolsters eIF4A-sensitive translation of selected host cell transcripts, including the one encoding the immunomodulatory cytokine TGF-β.

**Figure 5.**
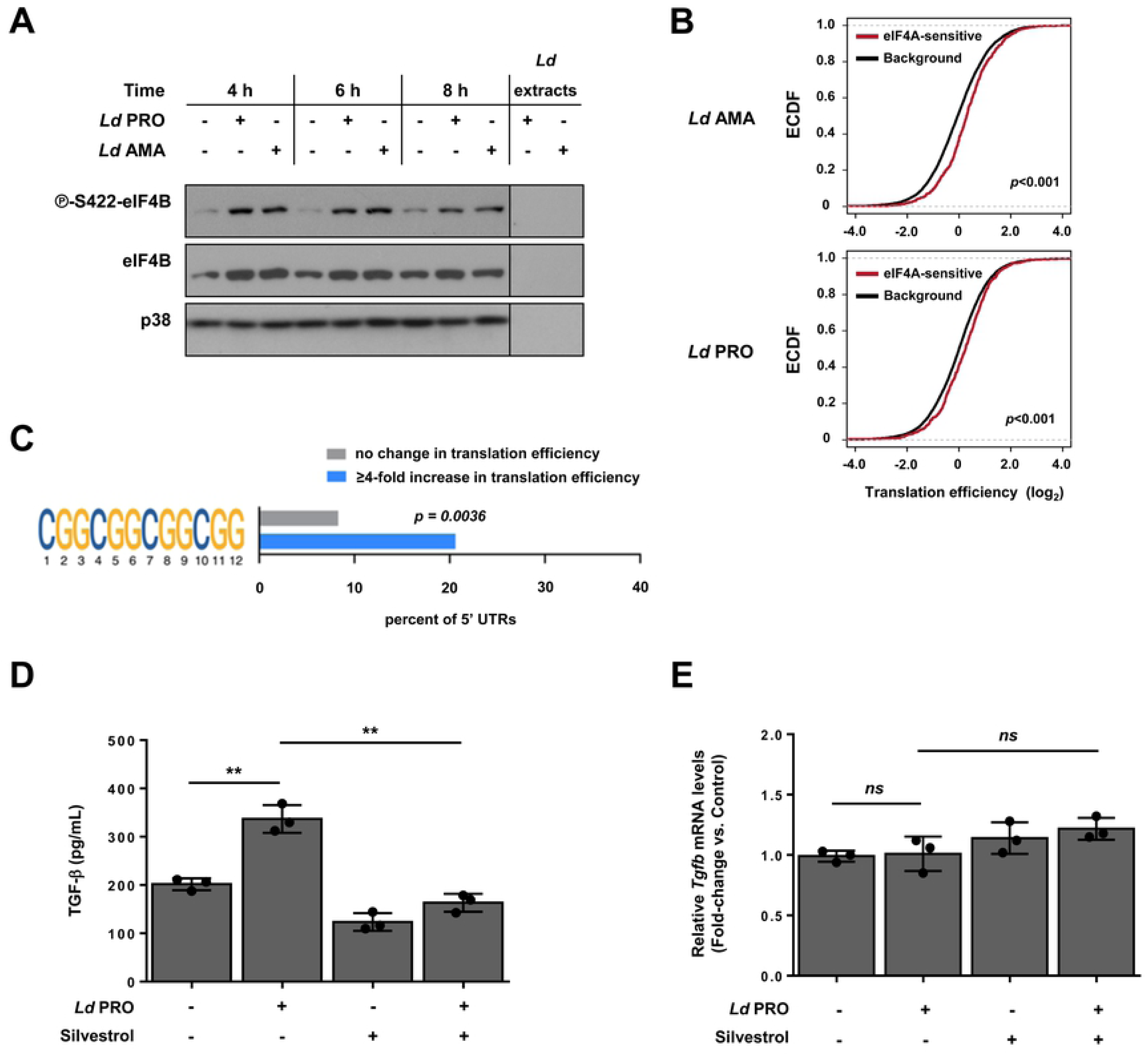
*L. donovani* infection activates eIF4A-sensitive mRNA translation in macrophages. (**A**) BMDM cultures were inoculated with *Ld* AMA, *Ld* PRO or left uninfected for the indicated times. Phosphorylation status and expression levels of eIF4B were monitored by Western blotting. Total levels of p38 MAPK were used as a loading control. (**B**) Empirical cumulative distribution function (ECDF) of translational efficiencies (infection vs. control) for the compilation of previously reported eIF4A-sensitive transcripts as compared to those of all detected transcripts (background). Differences in translational efficiencies between eIF4A-sensitive vs control transcripts were assessed per parasite stage. (**C**) Percentages of infection-associated translationally activated mRNAs (≥4-fold increase) with at least one (CGG)_4_ motif in their 5’ UTR as compared to a random set of unchanged mRNAs. Fisher’s exact test was used to compare frequencies of the (CGG)_4_ motif between the transcript subsets and the resulting *p*-value is indicated. (**D**-**E**) BMDM cultures were pre-treated with 25 nM silvestrol or vehicle for 2 h, then were inoculated with *Ld* PRO or left uninfected for 6 h. (**D**) Secreted levels of TGF-β were measured by sandwich ELISA. (**E**) Relative amount of *Tgfb1*mRNA (normalized to *Actb*) was measured by RT-qPCR. (**A**) Results are representative of three independent experiments. (**B-C**) Data analyses were performed on samples generated from at least three biological replicates. (**D**-**E**) Data are presented as mean ± SD (biological replicates, n=3). \**p* <0.05, \*\**p* <0.01 (for the indicated comparisons), *ns* = non-significant.

### *L. donovani* survival within macrophages is differentially modulated through mTOR and eIF4A activity

During the course of infectious diseases, translational control acts as a host defense mechanism but can also be exploited by the invading pathogen as a survival strategy (44). In regard to infections caused by protozoan parasites, augmented mTOR-sensitive translation was associated with parasite persistence during *Toxoplasma gondii* infection in macrophages (45) whereas eIF4A inhibition suppressed progression of cerebral malaria (12). These findings in combination with activation of mTOR- and eIF4A-sensitive translation in BMDMs upon parasite infection (**Fig 3** and **Fig 5**, respectively), prompted us to investigate the role of these translational regulators for *L. donovani* survival within the host cell. To address this, BMDMs were pre-treated with either rapamycin or silvestrol and infected with *L. donovani* promastigotes. Interestingly, parasite numbers increased in presence of rapamycin at 24 h post-infection (∼92% increase compared to DMSO control) (**Fig 6A**) whereas the opposite effect was observed upon cell exposure to silvestrol (∼57% reduction compared to DMSO control) (**Fig 6B**). These data indicate that mTOR limits *L. donovani* persistence within the host cell while eIF4A promotes it.

**Figure 6.**
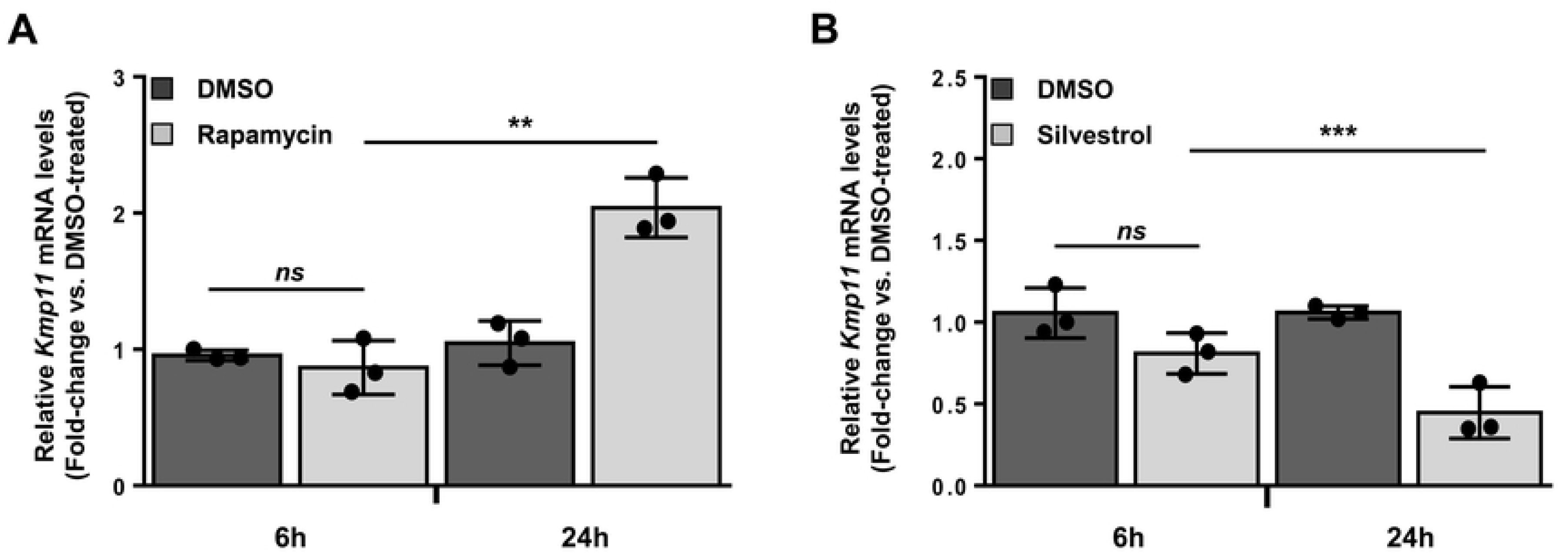
Host mTOR and eIF4A differentially regulate *L. donovani* persistence within macrophages. BMDM cultures were treated with (**A**) 20 nM rapamycin, (**B**) 25 nM silvestrol or an equivalent volume of DMSO (vehicle) for 2 h, then inoculated with *L. donovani* parasites (MOI 10) for 6 and 24 h. Quantification of intracellular parasites was performed by measuring the relative amount of *Leishmania Kmp11* mRNA (normalized to *Actb*) by RT-qPCR. Data are presented as mean ± SD (biological replicates, n=3). \*\**p* <0.01, \*\*\**p* <0.001 (for the indicated comparisons), *ns* = non-significant.

## Discussion

Transcriptome-wide analyses of mRNA translation in infected cells and tissues have revealed profound yet selective perturbation of the host translatome during infections caused by intracellular pathogens, including viruses and the protozoan parasite *Toxoplasma gondii* (45–47). *L. donovani* infection causes widespread changes in host gene expression (4–8). Yet, differences between the transcriptome and the proteome of infected cells suggested that post-transcriptional mechanisms may also contribute to establishing the post-infection proteome (9). Herein, using polysome-profiling, we demonstrate early translational reprogramming in macrophages following infection by *L. donovani* amastigotes and promastigotes. The majority of the changes in the translatome of the host cell were induced in a similar fashion by the two life stages of the parasite, including activation of both mTOR- and eIF4A-sensitive translation programs encoding central regulators of inflammation and mRNA metabolism. Unexpectedly, inhibition of mTOR promoted parasite survival within the host cell whereas inhibition of eIF4A activity had the opposite effect, suggesting that selective modulation of host mRNA translation plays a dual role during *L. donovani* infection in macrophages.

Previous high-throughput studies identified numerous biological processes affected in macrophages during *L. donovani* infection (5, 6, 8, 9). Translational profiling of infected macrophages indicates that in addition to altered mRNA levels, *L. donovani* selectively adjusts the proteome of the host cell to its own benefit by modulating the translational efficiencies of subsets of mRNAs. In line with dysregulated antigen presentation during *L. donovani* infection (48), transcripts encoding several MHC class I components were translationally suppressed in *L. donovani*-infected macrophages. In contrast, translation of mRNAs related to chromatin remodeling (i.e. histones and DNA- and histone-modifying enzymes) was augmented in cells infected with *L. donovani*. This suggests translational control of parasite-directed epigenetic changes known to inhibit innate immune responses of the host cell (49). Supporting this notion, a proteomics study identified several histones and chromatin remodelling proteins induced in macrophages during *L. donovani* infection which correlated with greater transcriptional activity (9).

Additional biological processes targeted during VL are related to host mRNA metabolism (i.e. stability, splicing and translation) (9, 50). Consistently, we identified numerous macrophage transcripts encoding translation initiation factors, splicing factors and RBPs with altered translational efficiencies during *L. donovani* infection. The mouse genome encodes more than a thousand of RBPs (51), including a subset we identified as translationally activated upon *L. donovani* infection. Regulation of RBPs is of particular interest during VL as they play a central role during immune responses (28). Further investigation is required to assess the impact of translational control of host RBP and mRNA metabolism for the outcome of *L. donovani* infections.

Our data showing that mTOR inhibition favors parasite survival within macrophages suggest that mTOR activation is part of a host defense mechanism against *L. donovani*. Further supporting this notion, phosphorylation of mTOR downstream targets S6K1 and 4E-BP1 occurred early during infection and rapidly decreased thereafter, an event that could be at least in part mediated by host phosphatases known to be activated upon *Leishmania spp.* infection (3). Upregulation of mTOR signaling in *L. donovani*-infected macrophages was paralleled by a significant increase in translational efficiencies of a large number of mTOR-sensitive transcripts characterized by the presence of a 5’ TOP motif. In particular, we showed that PABPC1, an RBP whose encoding mRNA harbors a TOP-motif (15), is induced during infection in an mTOR-dependent fashion. PABPC1 regulates stability and translation of mRNAs (52). Notably, PABPC1 is part of an inhibitory translational complex along with the zing finger protein 36 (Zfp36) that prevents overexpression of pro-inflammatory mediators in activated macrophages (29). Therefore, it is plausible that PABPC1 binds to specific host mRNAs to dampen macrophage inflammatory responses during *L. donovani* infection.

In addition to PABPC1, our data indicate that EIF2AK2 is upregulated in *L. donovani*-infected macrophages through mTOR-dependent mechanisms. Given that *L. donovani* triggers TLR signaling in macrophages (35) and that both mTOR signaling and EIF2AK2 expression are augmented in response to TLR stimulation (34, 53), TLR-dependent mTOR activation might account for increased EIF2AK2 synthesis in *L. donovani*-infected macrophages. Accumulating evidence suggests that the role of EIF2AK2 during *Leishmania spp* infection varies between parasite species. For example, EIF2AK2 activity is induced during *L. amazonensis* infection and promotes parasite survival within macrophages through induction of an Nrf2-dependent antioxidant response (30, 54). Conversely, *L. major* prevents the activation of EIF2AK2 to avoid parasite clearance via EIF2AK2-inducible TNF production (30, 31). We observed that eIF2α phosphorylation decreases upon infection despite upregulation in EIF2AK2 expression and phosphorylation. This may be explained by the concomitant increase in mTOR activity which can recruit NCK1 to eIF2β and thereby reduce eIF2α phosphorylation to bolster ternary complex formation (55). Thus, the relative activity of eIF2α kinases and phosphatases may determine eIF2α phosphorylation-status and thereby downstream effects on gene expression and biology. Herein, regardless of the precise mechanism, the increased level of EIF2AK2 does not appear activate the integrated stress response (56).

Early phosphorylation of eIF4B and enrichment of reported eIF4A-sensitive mRNAs among translationally activated transcripts support that eIF4A also affects the translatome of *L. donovani*-infected macrophages. Consistently, silvestrol-mediated inhibition of eIF4A dampened *L. donovani* survival within macrophages while no effect was observed in extracellular parasite viability. The activity of eIF4A facilitates translation of transcripts encoding immune modulators (e.g. CXCL10, IRF1, IRF7, iNOS, STAT3, TGF-β, etc.) in various cell types (18, 20, 39, 40, 57) including macrophages (12, 22). Similarly, we observed that translation of eIF4A-sensitive transcript *Tgfb* and subsequent TGF-β production were upregulated in *L. donovani*-infected in macrophages. Of note, TGF-β is a cytokine associated with immune suppression and resistance to treatment during human and experimental VL (42, 43). In view of these previous studies and our present findings, it is conceivable that eIF4A-sensitive translation contributes to the immuno-pathogenesis of VL. Further characterization of eIF4A translational outputs in *L. donovani* infected cells and tissues will shed light on this matter.

The outcome of VL is defined by a complex network of converging molecular events at the interface between the parasite and the host (1). Herein, we have uncovered vast reprogrammed translation of protein-coding mRNAs expressed in macrophages upon infection by *L. donovani* amastigotes or promastigotes. Accordingly, numerous host transcripts critical during infection, including regulators of mRNA metabolism and inflammation, were found to be under translational control. Notably, our data indicate that some of these changes contribute to parasite clearance whereas others favor parasite persistence within the host cell, hinting at the therapeutic potential of perturbing specific host translation programs to control the infection.

## Materials and Methods

### Reagents

Culture media and supplements were purchased from Wisent, Gibco, and Sigma-Aldrich; rapamycin was obtained from LC Laboratories; torin-1 was provided by Cayman; 10-phenanthroline monohydrate was acquired from Sigma-Aldrich; silvestrol was purchased from Biovision.

### Parasites

*L. donovani* (LV9 strain) amastigotes were isolated from the spleen of infected female Golden Syrian hamsters (Harlan Laboratories) as previously described (58). *L. donovani* (LV9 strain) promastigotes were differentiated from freshly isolated amastigotes and were cultured at 26°C in M199 medium supplemented with 10% heat-inactivated FBS, 100 µM hypoxanthine, 5 µM hemin, 3 µM biopterin, 1 µM biotin, 100 U/mL penicillin, and 100 μg/mL streptomycin. Early passage stationary phase promastigotes were used for macrophage infections.

### Ethics Statement

Housing and experiments were carried out under protocols approved by the Comité Institutionnel de Protection des Animaux (CIPA) of the INRS – Centre Armand-Frappier Santé Biotechnologie (CIPA 1308-04 and 1710-02). These protocols respect procedures on good animal practice provided by the Canadian Council on animal care.

### Differentiation of bone marrow-derived macrophages

Bone marrow-derived macrophages (BMDM) were generated from precursor cells from murine bone marrow, as previously described (45). Briefly, marrow was extracted from bones of the hind legs, red blood cells were lysed, and progenitor cells were resuspended in BMDM culture medium supplemented with 15% L929 fibroblast-conditioned culture medium (LCCM). Non-adherent cells were collected the following day and were cultured for 7 days in BMDM culture medium supplemented with 30% LCCM with fresh medium replenishment at day 3 of incubation.

### Infection of bone marrow-derived macrophages

BMDM cultures were inoculated with *L. donovani* promastigotes or amastigotes at a multiplicity of infection (MOI) of 10:1, as previously described (59). Prior to infection, cells were serum-starved for 2 h and treated with inhibitors, when applicable.

### Viability assays

Viability of BMDM and extracellular *L. donovani* parasites was determined by the resazurin assay as described (11). Briefly, cells were treated with increasing concentrations of rapamycin (2.5 – 160 nM), torin-1 (12.5 – 800 nM), silvestrol (0.8 – 100 nM) or an equivalent volume of DMSO (vehicle) for 24 h at 37°C, 5% CO_2_. Medium was removed and replaced with fresh culture medium supplemented with 0.025% resazurin. Cultures were incubated for 4 h in presence of the inhibitors or DMSO at 37°C, 5% CO_2_. Optical density was measured using a Multiskan GO (Thermo-Fisher) at 600 and 570 nm. Absorbance at 600 nm was subtracted from readings at 570 nm. Experiments were performed in biological replicates (*n* =2); each sample was analyzed in a technical triplicate, the average of which was plotted against increasing concentrations of the respective inhibitor.

### Quantification of intracellular parasites

BMDM cultures were treated with 20 nM rapamycin, 25 nM silvestrol or an equivalent volume of DMSO (vehicle) for 2 h, then inoculated with *L. donovani* parasites (MOI 10:1) for 6 and 24 h. Relative quantification of intracellular parasites was performed by RT-qPCR measuring the expression of *Leishmania kmp11* gene as described (60). Primer sequences are listed in **S4 Table**.

### Quantitative RT-PCR

Purified RNA (500 ng) was reverse transcribed using the Superscript IV VILO Master Mix (Invitrogen). Quantitative PCR was performed with PowerUp™ SYBR^®^ Green Master Mix (Applied Biosystems). Relative quantification was calculated using the comparative Ct method (ΔΔCt) (61) and relative expression was normalized to mouse *Actb*. Experiments were performed in independent biological replicates (n=3); each sample was analyzed in a technical triplicate, the average of which was plotted against the respective conditions used. Primers were designed using NCBI Primer-BLAST (http://www.ncbi.nlm.nih.gov/tools/primer-blast/) (**S4 Table**).

### Western blot analysis

Following infection and other treatments, total cell lysates were prepared for SDS-PAGE and Western blotting as described (59). Primary antibodies anti-mTOR (#2983), anti-phospho-S6K1 (T389; #9234), anti-phospho-RPS6 (S235/236; #2211), anti-phospho-RPS6 (S240/244; #5364), anti-phospho-eIF4B (S422, #3591), anti-eIF4B (#3591), anti-phospho-4E-BP1 (T37/46; #2855), anti-4E-BP1 (#9644), anti-phospho-eIF2α (S51; #3398), anti-eIF2α (#2103), anti-PABCP1 (#4992), and β-actin (#3700) were purchased from Cell Signaling Technologies; anti-phospho-PKR (EIF2AK2) (T451; #07-886) was obtained from Millipore; anti-RPS6 (#sc-74459), anti-S6K1 (#sc-230), and anti-PKR (EIF2AK2) (#sc-6282) were acquired from Santa Cruz Biotechnology. Horseradish peroxidase (HRP)-linked goat anti-rabbit and goat anti-mouse IgG secondary antibodies were purchased from Sigma-Aldrich.

### RNA fractionation and purification from polysome fractions

Cytosolic lysates of infected and control BMDM were prepared for RNA fractionation as described (45). Lysates were layered over 5 to 50% sucrose density gradients and sedimented using a Beckman SW41 rotor at 36,000 rpm (= 221,830.9 × *g*) for 2 h at 4°C. Gradients were fractionated and collected (30 s, 500 µL/fraction), and the absorbance at 254 nm was recorded continuously using a Brandel BR-188 density gradient fractionation system. Input material (total cytoplasmic mRNA) and efficiently translated mRNA (heavy polysome-associated, >3 ribosomes) were extracted with QIAzol (Qiagen) and purified using RNeasy MinElute Cleanup Kit (Qiagen). Purity and integrity of RNA was assessed using a Bioanalyzer 2100 with a Eukaryote Total RNA Nano chip (Agilent Technologies).

### RNAseq data processing

RNAseq libraries were generated using the Smart-seq2 method (62) to enable polysome-profiling of small samples as described previously (63). Libraries were sequenced by using an Illumina HiSeq2500 instrument with a single-end 51-base sequencing setup for both total cytoplasmic and polysome-associated mRNAs (mRNAs associated with >3 ribosomes) from three independent biological replicates for uninfected and *L. donovani* promastigote-infected BMDM, and five independent biological replicates for *L. donovani* amastigote-infected BMDM. First, RNAseq reads mapping to the reference genome of the Nepalese BPK282A1 strain of *L. donovani* were removed. The filtered reads were then aligned to the mouse genome mm10. HISAT2 was used for all alignments with default settings (64). Gene expression was quantified using the RPKMforgenes.py script (http://sandberg.cmb.ki.se/media/data/rnaseq/rpkmforgenes.py) (62) with options -fulltranscript -readcount -onlycoding from which raw per gene RNAseq counts were obtained (version last modified 07.02.2014). Genes that had zero counts in all samples were discarded. Annotation of genes was obtained from RefSeq.

### RNAseq data analysis using anota2seq

RNAseq counts were normalized within anota2seq using default TMM-log2 normalization (23). Significant changes in translation, abundance and buffering were identified by anota2seq (23) using default parameters with the following modifications: minSlopeTranslation=-0.5; maxSlopeTranslation=1.5; FDR ≤0.15; *apvEff* >log_2_(2.0); deltaPT>log_2_(1.25); and deltaP>log_2_(1.5). In anota2seq model one, infections were compared to control (i.e. *Ld* PRO versus control and *Ld* AMA versus control); in model two, cells infected by different parasite stages were compared together with a contrast to complete the anota2seq model (i.e. *Ld* PRO vs *Ld* AMA and *Ld* PRO versus control). Identifiers for genes which cannot be distinguished based on their high sequence similarity (also reported by RPKMforgenes.py), were excluded from downstream analyses. For further analysis, we used a published list of TOP-containing mRNAs (27) and previously reported translational signatures of eIF4A-sensitive mRNAs (36, 38–40). The difference in log_2_ fold-change of translational efficiency (i.e. *apvEff*) between the signatures and the background was assessed using Wilcoxon-Mann-Whitney tests.

### Gene ontology analyses

Gene ontology analyses were performed using the panther tool (25) of the Gene Ontology Consortium (http://geneontology.org/) on the union of transcripts activated or inhibited in BMDMs infected by *L. donovani* amastigotes or promastigotes. Heatmaps of translational efficiencies of transcripts activated or inhibited in BMDMs infected by *L. donovani* amastigotes or promastigotes were generated using Morpheus. (https://software.broadinstitute.org/morpheus/index.html, Broad Institute)

### RNA sequence motif analyses

Non-redundant RefSeq 5’ UTRs were retrieved from genome build mm10 using the UCSC Table Browser (https://genome.ucsc.edu/cgi-bin/hgTables). Analysis of Motif Enrichment (AME) was performed on the RefSeq-annotated 5’ UTR sequences from transcripts with ≥4-fold increases in TE in *L. donovani*-infected BMDM (402 5’ UTRs) compared to a control list of 228 randomly selected 5’ UTRs from the set of non-translationally regulated transcripts. The presence of the previously described guanine quartet (CGG)_4_ motif (39) in both lists was scored and a one-tailed Fisher’s exact test was performed to determine significance of enrichment.

### Statistical Analysis

Where applicable, data are presented as the mean ± standard deviation (SD) of the mean. Statistical significance was determined by using one-way ANOVA followed by a Tukey *post-hoc* test; calculations were performed by using Prism 7 software package (GraphPad). Differences were considered significant when \**p* < 0.05, ** *p* < 0.01, *** *p* < 0.001.

## Acknowledgements

The authors thank the Science for Life Laboratory, the National Genomics Infrastructure, NGI, and Uppmax for providing assistance in massive parallel sequencing and computational infrastructure.

## Author Contributions

Conceived and designed the experiments: VC, LPL, LM, AZ, OL, MJ. Performed the experiments: VC, LPL, LM AZ, GAD. Analyzed data: VC, LPL, LM, JL, TEG, AZ, TA, OL, MJ. Contributed materials, methods, and/or technology: GAD, AD, TA, OL. Wrote the manuscript: VC, LPL, OL, MJ.

## Data availability Statement

All relevant data are within the paper.

## Supporting information captions

**S1 Figure. Differential regulation of host mRNA translation between *L. donovani* promastigotes and amastigotes.** (**A**) Kernel densities of adjusted *p* values (FDRs) following anota2seq analysis on changes in total mRNA, heavy polysome-associated mRNA, translational efficiency, and translational buffering comparing *Ld* PRO to *Ld* AMA-infected BMDM (n ≥ 3). (B) Scatter plot of log_2_ fold changes (for the same comparisons as in panel A) for heavy polysome-associated mRNA and total cytosolic mRNA. Differentially regulated transcripts through translation, abundance or buffering are indicated. Unchanged mRNAs are shown in grey (n ≥ 3) (**C**) Venn diagrams indicating the number of mRNAs up- or down-regulated at the level of translation, abundance, and buffering for *Ld* PRO-infected BMDM compared to *Ld* AMA-infected BMDM.

**S2 Figure. *L. donovani* infection promotes EIF2AK2 but not eIF2α phosphorylation.** BMDM cultures were inoculated with *Ld* AMA, *Ld* PRO or left uninfected for 6 h. Phosphorylation and expression levels of EIF2AK2 and eIF2α were monitored by Western blotting. Total amounts of β-actin were used as a loading control. Total protein extracts from *Ld* cultures were used to control for any cross-reactivity of the antibodies against parasite antigens. Data are representative of three independent experiments.

**S3 Figure. Measurement of acute toxicity of mTOR inhibitors on extracellular *L. donovani* promastigotes.** *L. donovani* cultures were treated with increasing concentrations of rapamycin (2.5 – 160 nM), torin-1 (12.5 – 800 nM) or an equivalent volume of DMSO (vehicle) for 24 h. Acute toxicity of the inhibitors was measured by resazurin assays. Percent viability was normalized to DMSO-treated parasites. Data are representative of two independent experiments performed in technical triplicates.

**S4 Figure. Measurement of acute toxicity of silvestrol on BMDMs and *L. donovani* promastigotes.** (**A**) BMDM and (**B**) *L. donovani* cultures were treated with increasing concentrations of silvestrol (0.8 – 100 nM) or an equivalent volume of DMSO (vehicle) for 24 h. Acute toxicity of the inhibitor was measured by resazurin assays. Percent viability was normalized to DMSO-treated parasites. Data are representative of two independent experiments performed in technical triplicates.

**S1 Table. List of host mRNAs differentially regulated upon *L. donovan*i infection.** Anota2seq identified 1516 host mRNAs upregulated (first tab) and 1592 downregulated (second tab) via changes in translational efficiency. The abundance of 1236 (third tab) and 914 (fourth tab) host transcripts was found to be increased or decreased, respectively; 369 (fifth tab) and 258 (sixth tab) host mRNAs were translationally buffered after showing increased or decreased abundance, respectively; 137 host transcripts were found to be differentially regulated through translation, 765 were regulated via changes in abundance, and 71 through buffering between amastigote-infected and promastigote-infected BMDMs. Fold-change (log2) and FDR values are listed for each target and stage.

**S2 Table. Gene Ontology enriched categories for translationally-regulated mRNAs upon *L. donovani* infection.** The *Panther* tool of the Gene Ontology (GO) Consortium identified enriched categories by *Biological Process* (first tab), *Molecular Function* (second tab) and *Cellular Component* (third tab) on host mRNAs translationally regulated upon *L. donovani* infection. FDR values, fold enrichment and number of transcripts per category are listed for significantly enriched categories (FDR <0.05, mRNAs ≥5).

**S3 Table. List of TOP, eIF4A-sensitive, and (CGG)_4_ motif-harboring transcripts found in the translationally regulated dataset of *L. donovani*-infected BMDMs.** TOP mRNAs (27) (first tab), eIF4A-sensitive mRNAs (36, 38–40) (second tab), and (CGG)_4_ motif-harboring mRNAs (39) (third tab) found in the translationally up-regulated dataset of *L. donovani*-infected BMDM.

**S4 Table. Primer sequences used for RT-qPCR analyses.**

